# Human Stem Cell-derived Kidney Collecting Duct Model via Epithelial Microphysiological Analysis Platform: Epi-MAP

**DOI:** 10.1101/2024.11.11.620553

**Authors:** SoonGweon Hong, Minsun Song, Ankit Patel, Kyle W. McCracken, Joseph V. Bonventre, Luke P. Lee

## Abstract

The kidney epithelium’s pivotal role in molecular filtration, metabolism, and excretion highlights the crucial importance of understanding kidney physiology in drug development. However, our knowledge is largely derived from non-human or non-physiological models, potentially limiting its applicability to humans. To address this significant gap, we have pioneered a human kidney epithelial microphysiological analysis platform (Epi-MAP) designed to establish, mature, and monitor renal functions of the human collecting epithelium within a physiologically relevant microenvironment. We first demonstrate the highly mature collecting duct physiology derived from human stem cells, enabled by the Epi-MAP’s microenvironments that recapitulate in vivo asymmetries in fluidic and biochemical conditions. The integrated biosensors of the Epi-MAP provide long-term, time-resolved epithelial maturation trajectories, revealing advanced integrity and functional maturity with transepithelial metrics. Furthermore, Epi-MAP’s electrophysiological analytics for measuring water flux, in conjunction with transepithelial potential and resistance, allow for real-time decoding of intricate epithelial responses to substance stimulation, showcasing its effectiveness as a robust pharmacological test model. This human cell-derived, physiologically advanced model on a chip stands as a robust in vitro tool, offering comprehensive insights into human kidney biology and significantly enhancing drug discovery process based on human physiology.

## INTRODUCTION

While animal models have fueled kidney research, providing valuable insights into structure and function^1-5^, interspecies differences arising from factors like evolutionary pressure due to size, diet, and habitat can lead to inaccurate evaluations of physiological dynamics in the human kidney^6-8^. This discrepancy frequently results in misjudgments of safety and efficacy during drug discovery^9-13^. Additionally, the disparity in molecular factors and pathogenesis with the human kidney renders animal models non-predictive for the development of effective therapies for human kidney diseases^14-17^. Recognizing these limitations, researchers are increasingly seeking to develop more physiologically relevant human kidney models to advance our understanding of renal biology and improve precision in drug discovery.

Human kidney biology remains challenging, largely due to the scarcity of functional, physiologically relevant human cell models. Unlike other organs, the human kidney has yielded few immortalized cell lines capable of replicating full-fledged functions, even within limited contexts. Even fresh primary cells, while offering glimpses into native biology, face challenges such as ethical concerns, biopsy difficulties, lineage instability, rapid functional decline, and inter-individual variability^18,19^. Consequently, the development of in vitro kidney models that accurately mirror intricate functions and allow controllable, reproducible studies has been elusive. This scarcity of suitable models has significantly impeded our comprehensive exploration of human kidney biology.

A renaissance is dawning in kidney research, driven by breakthroughs in ex vivo differentiation techniques, particularly pluripotent stem cell-derived kidney organoids^20-23^. These miniature, self-organizing structures hold immense promise for unlocking the intricacies of human kidney development and offer an effective method to generate diverse kidney cell types otherwise challenging to obtain through direct differentiation. Our recent work exemplifies this potential^24^. We successfully generated human stem cell-derived ureteric bud organoids and differentiated them into functional renal collecting duct (hCD) cells. These cells exhibit basic CD functions like tight junction formation and ion transport. However, the study lacked dynamic physical and biochemical cues across both time and space domains, which are crucial for capturing the full spectrum of hCD functionalities and their responses to external stimuli. This limits our ability to fully utilize them for studying human kidney physiology.

To unlock the full potential of the hCD cells, we developed our epithelial microphysiological analysis platform (Epi-MAP), a meticulously designed kidney epithelial microfluidic platform integrated with in-situ sensors. Epi-MAP fosters the development of the early-stage kidney cells to highly functional hCD epithelia by providing physiologically relevant physiochemical dynamics. The fidelity and precision of Epi-MAP’s long-term culture enable the reproducible induction of key physiological functions including transport elements, epithelial integrity, and signaling pathways. Notably, Epi-MAP’s groundbreaking in situ electrophysiology unveils the hidden orchestra of hCD function, capturing intricate dynamics from fleeting seconds to sustained month-long adaptations, offering unprecedented insights into its diverse functions and unlocking a new era of kidney physiology understanding. This pioneering human kidney model, enabled by a reproducible human cell source and bioengineering recapitulation of physiological microenvironments, holds great potential to revolutionize our understanding of renal physiology and pave the way for the development of more accurate and effective therapeutic strategies for a wide range of kidney diseases.

## RESULTS

### Kidney Epi-MAP Emulates and Monitors Physiological Contexts of Kidney Tubules

Mirroring the native kidney’s tubular architecture defining the lumen and the interstitium, our kidney Epi-MAP (Fig. 1) features two microfluidic channels separated by a nanoporous membrane mimicking the epithelial basement membrane. Upon reaching confluency, the apical surface of the epithelial monolayer is subjected to a precisely controlled laminar flow in the top channel, replicating urinary tubular flow with physiologically relevant shear stress (∼0.2 dyn·cm^-2^)^25^. The bottom channel simulates interstitial fluid surrounding the hCD with a controlled flow rate of 0.1 mL·min^-1^. The low-profile design (30 μm height) of the basolateral channel ensures that its biochemical composition dynamically adapts to epithelial transport, enabling well-reflected readouts by our suite of real-time microfluidic sensors.

**Figure 1.**
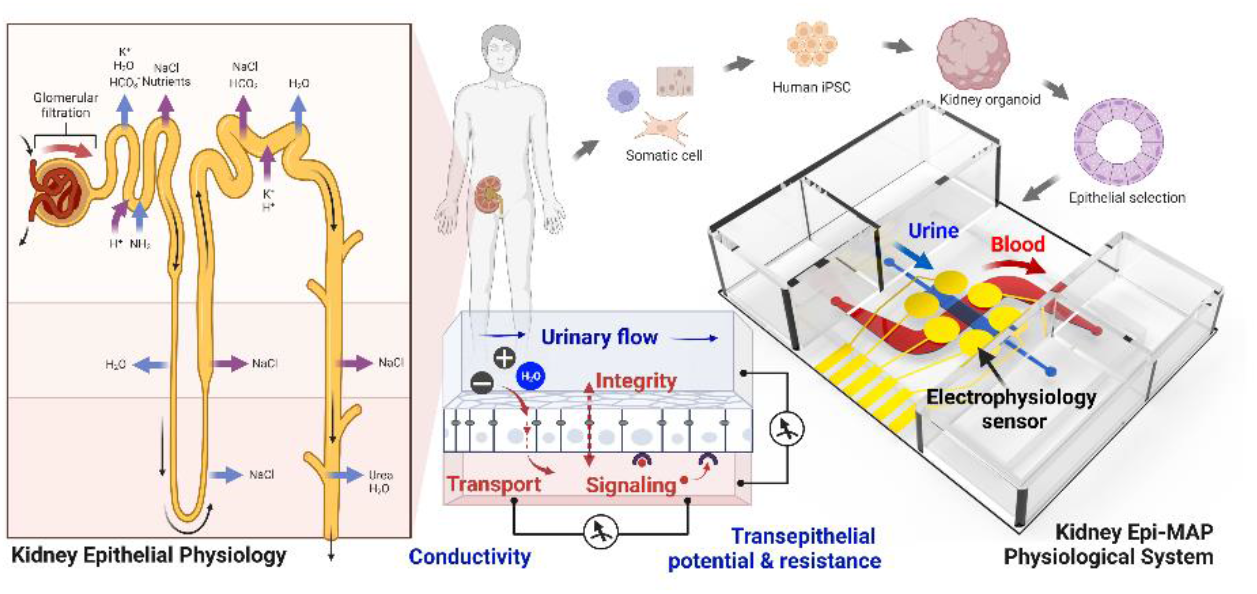
Recapitulating Human Collecting Duct Physiology in the Epithelial Microphysiological Analysis Platform (Epi-MAP). This study recreates human collecting duct epithelial physiology utilizing collecting duct cells derived from human stem cells within the integrated microphysiological system, Epi-MAP. Employing stem cell and organoid techniques, we establish a dedicated “humanized” in vitro resource of collecting duct-specific epithelial cells capable of proliferation and potential physiological expression. Epi-MAP, a microfluidic platform integrated with biosensors, generates in vivo-like fluid flow and biochemical gradients, allowing for precise long-term functional maturation of kidney epithelial function. Moreover, Epi-MAP enables real-time, multimodal monitoring of epithelial biology directly within the system. This powerful combination positions Epi-MAP as a promising human model, advancing our understanding of collecting duct biology and its role in health and disease.

Our central hypothesis posits that long-term maturation of hCD cells within the Epi-MAP system can promote robust development of epithelial functions comparable to their in vivo counterparts. Mimicking the natural process of neonatal kidney tubules that acquire physiological maturation through gradual exposure to dynamic flow and biochemical gradients^26^, we designed a long-term, biochemically staged, dynamic culture in Epi-MAP. After establishing a confluent epithelial monolayer in conventional media that promotes proliferation and defining distinct luminal and interstitial compartments, the cells undergo months of maturation in an in vivo-mimetic biochemical microenvironment. To enable a controlled developmental trajectory, we implement a system providing biochemically and fluidically consistent unidirectional flow through large media reservoirs and a computer-operated tilting plate. This eliminates the need for cumbersome tubing and pumping in microfluidic operation, thus enhancing experimental accuracy, reproducibility, and enabling high-throughput studies with minimal operational interventions. In our optimal setup, the long-term precision of fluidics and biochemical conditions is demonstrated to enhance the reproducibility of hCD physiology, in tandem with a consistent human stem cell-derived cell source, within the Epi-MAP system.

Epi-MAP’s innovative design enables unprecedented in situ, real-time electrophysiological monitoring within a standard cell culture incubator. Six strategically positioned surface electrodes, nestled within dedicated via chambers adjacent to fluidic channels, seamlessly capture cellular activity without disrupting fluid dynamics. This configuration, connected to a remotely located electrochemical analyzer via a programmed multiplexing switch, facilitates diverse physiological measurements. The microfabricated, electrochemically stable platinum electrodes operate in pairs, empowering long-term impedance and open circuit potential measurements in multiple configurations. These measurements reveal transepithelial physiological attributes, including epithelial integrity and vectorial ion transport, by assessing transepithelial potential difference (TEPD), transepithelial electrical resistance (TEER), and interstitial fluid conductivity. This realtime, in-situ monitoring capability permits precise and continuous assessment of epithelial functions and physiological dynamics in Epi-MAP, offering comprehensive insights into kidney physiology and unlocking a new era of kidney research.

Beyond its innovative platform design, Epi-MAP incorporates spatially distinct biochemical environments to mimic the in vivo fluid complexity of the tubular epithelium. Leveraging insights from single-cell transcriptomic data^24^, we tailored the luminal and interstitial media to reflect the specific needs of our inner medullary hCD cells. The luminal fluid emulates urine composition with elevated sodium and urea, mimicking the conditions for these cells to perform their function in the fine-tuning of urine. In contrast, the interstitial compartment resembles the nutrient-rich, hyperosmotic inner medullary interstitium through the addition of these solutes to mirror the in vivo gradients. This carefully crafted configuration establishes a modest osmolarity gradient (20 mOsm·L^-1^) between the luminal (460 mOsm·L^-1^) and interstitial (480 mOsm·L^-1^) fluids, further enhancing the physiological realism of Epi-MAP (detailed further in Methods).

### Long-Term Culture in Epi-MAP Promotes Advanced Physiological Maturation of hCD Cells

To systematically investigate physiological effects of the microenvironmental aspects involved in our hCD modeling, we conducted a comparative examination of hCD models maintained in four culture methods, depicted in Fig. 2a. The first model was a conventional 2D culture on a dish, devoid of apicobasal polarity. The second model was a transwell culture designed to facilitate apicobasal transport, which has been shown to promote epithelial integrity and polarization^27^. The third model incorporated physiological media that mimics the biochemical microenvironment of the inner medullary hCD within the same transwell setup as the second model. Finally, the fourth model, Epi-MAP provided a dynamic culture environment with physiological media, allowing continuous delivery and removal of urine components, nutrients, and cell byproducts along multi-compartmentalized flows. Remarkably, as the physiological complexity increased across these models, we observed increasingly refined epithelial morphology, gene expression, and functional properties of the hCD cells.

**Figure 2.**
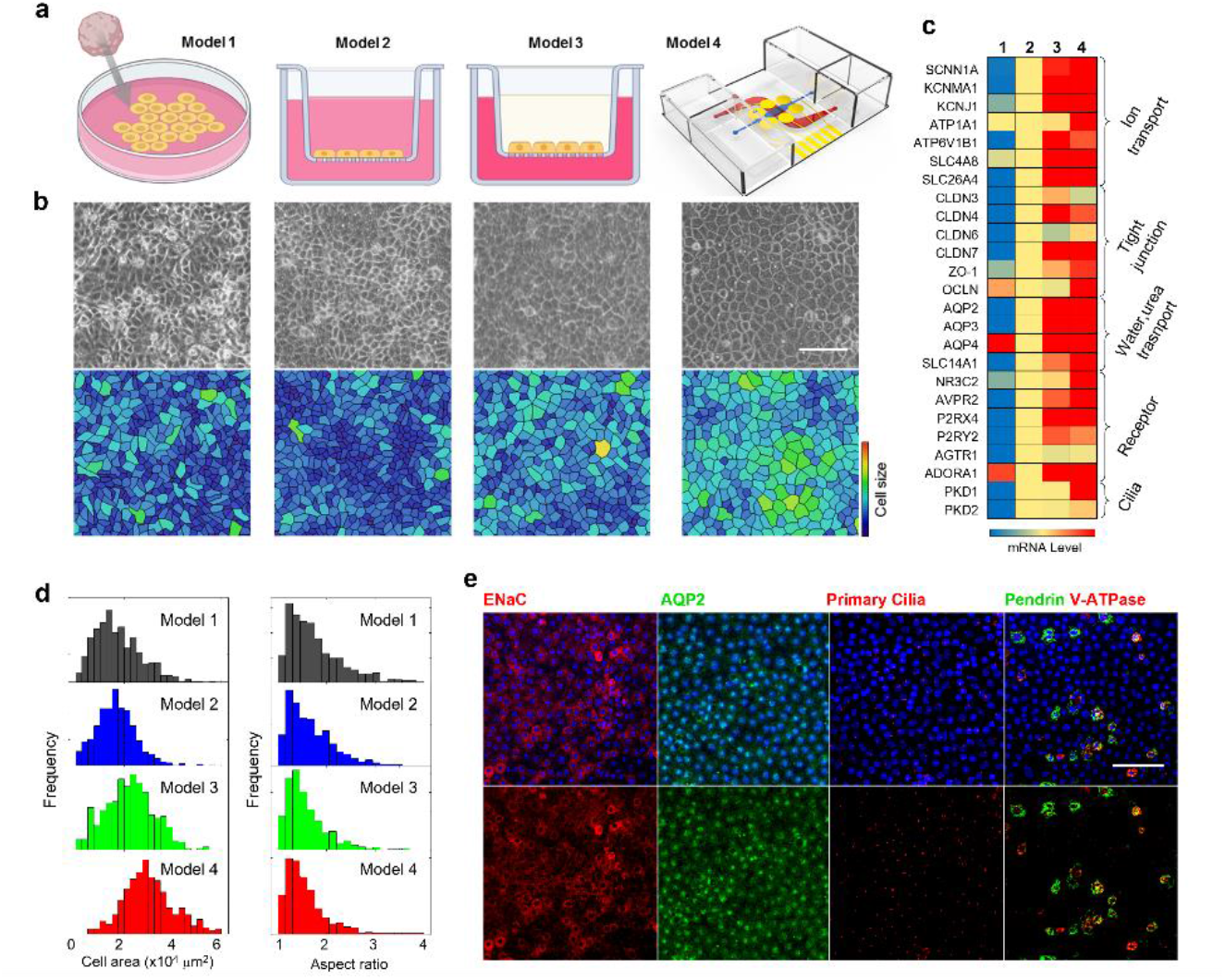
Advanced hCD Physiological Expression in Epi-MAP: Multiparameter Analysis Reveals Enhanced Gene, Protein, and Morphology. **a**, A four-model approach to explore microenvironment effects on human collecting duct physiology. The models progress in complexity: dish culture with basic media, transwell culture for polarity, physiological media transwell for biochemical cues, and finally, Epi-MAP combining all elements with fluid dynamics. **b**, Representative photographs and pseudo-images of the models, depicting enhanced morphology as microenvironment complexity increases. Scale bar: 50µm. **c**, Differential gene expression across models: Heatmap comparing mRNA levels of genes related to ion transport, tight junction, water/urea transport, signaling receptor, and cilia formation. **d**, Cell size and aspect ratio histograms for each model. Larger and more symmetrical cells appear in cultures as model complexity and physiological relevance increase. **e**, Immunostaining of the hCD epithelium in Epi-MAP for principal cell markers: epithelial sodium channel (ENaC), aquaporin 2 (AQP2), and primary cilia (stained with acetyl-α-tubulin) and intercalated cell markers: pendrin and V-ATPase. Blue overlays show DAPI nucleus staining (scale bar: 50µm). Ciliary expression under fluidic shear stress is further characterized in Supplementary Fig. 1.

Microscopic analysis of the four hCD models revealed starkly different epithelial monolayer morphologies (Fig. 2b). Lacking apicobasal polarity, Model 1 displayed a disorganized monolayer with localized areas of cell stacking after confluency. In contrast, Model 2 with enhanced apicobasal polarity, displayed an improved monolayer morphology with a more organized polygonal shape. The addition of physiological media and fluidic shear stress further enhanced epithelial morphology, resulting in increased cell size, more regular cell shape, and well-defined cellular borders (Fig. 2d). As expected for mature epithelia, morphological remodeling or cellular proliferation was rarely observed within the Epi-MAP culture, likely due to improved cell-cell adhesion and advanced maturation. These findings underscore the critical roles of apicobasal polarity, physiological media, and fluidic shear stress in maintaining a stable, organized hCD monolayer.

Immunostaining at day 60 in Epi-MAP culture unveiled a remarkable feature: a diverse hCD population mirroring the in vivo collecting duct, alongside confirmation of mature hCD characteristics (Fig. 2e). Consistent with our earlier finding^24^, most of the population showed characteristics of principal cells (PCs) with expression for ENaC and AQP2, as PCs are the major hCD population responsible for sodium and water regulation. Primary cilia, a crucial mechanosensor of PC, were similarly abundant in frequency to those observed in vivo, unlike in the static culture conditions of Model 3 (Supplementary Fig. 1). Notably, interspersed within the PC population was a small subset of cells expressing pendrin and V-ATPase, hallmark proteins of intercalated cells (ICs). This arrangement mirrors the in vivo collecting duct, where ICs, despite their limited numbers, play crucial roles in regulating ion and pH homeostasis, working in concert with PCs. The rare presence of ICs in the static culture (Model 3) and the absence of IC progenitors in single-cell RNA-seq of the original cell pool suggest PC-IC transdifferentiation by the dynamic environment of Epi-MAP. Further investigations are warranted to elucidate the precise mechanisms behind IC appearance; however, the mixed hCD population in Epi-MAP offers a potentially more physiologically advanced model for studying human kidney functions and modeling diseases.

Examining essential functional genes through RT-qPCR unveiled upregulation patterns in numerous genes associated with membrane ion transport, tight junctions, cilia formation, signaling receptors, and water/urea transport. These patterns indicate a progressive physiological advancement across the models (Fig. 2c). While each microenvironmental aspect contributed unique expression signatures, the Epi-MAP hCD model was particularly noteworthy for its highly consistent upregulation of genes crucial for collecting duct functionality.

Among the ion transport genes, the Epi-MAP model displayed significantly upregulated expression of the epithelial sodium channel (ENaC) subunit *SCNN1A* and Na^+^-K^+^ ATPase (*ATP1A1*), indicating enhanced sodium and potassium handling capacity. Conversely, the expression of the potassium channels BK (*KCNMA1*) and ROMK (*KCNJ1*) appeared more markedly affected by the presence of physiological media. Notably, the Epi-MAP model exhibited the highest expression of genes characteristic of B-type IC, including *SLC4A8* for the sodium-driven sodium bicarbonate exchanger NDCBE (facilitating bicarbonate reabsorption) and *SLC26A4* for pendrin (involved in chloride/bicarbonate exchange). This differential pattern suggests Epi-MAP promotes both PC and B-type IC functions.

Other genes followed similar trends, with peak expression in either Model 3 or 4. Notably, mRNA levels of *AQP2* (apical water channel), *SLC14A1* (urea transporter, UT-B), *NR3C2* (mineralocorticoid receptor), and *AVPR2* (vasopressin receptor), reached their highest abundance in the Epi-MAP model. Some genes showed greater responsiveness to apicobasal polarity, including those encoding tight junction proteins (*CLDN3, 4, 6*, and *7*) and receptors (*P2RY2* for purinergic receptor and *AGTR1* for angiotensin II receptor). The overall gene expression patterns suggest that the dynamic microenvironment of Model 4 effectively upregulates genes crucial for CD epithelium functions like transepithelial transport, tight junction integrity, and hormone responses. This confirms the potential of the Epi-MAP model to offer a more accurate and physiologically advanced model for studying human kidney function and disease.

### Physiologically Relevant Microenvironment of Epi-MAP Enhances Functional Maturation of Human Collecting Duct Epithelium

Prolonged in-situ electrophysiological monitoring within Epi-MAP revealed a profound influence of physiologically relevant conditions on the functional development of hCD, particularly their pivotal role in sculpting transepithelial properties as elucidated by TEPD and TEER metrics. First, we implemented a two-phased, long-term culture (both exceeding 20 days) with distinct biochemical compositions to elucidate the influence of in vivo biochemical environments. Phase 1, employing conventional epithelial media, facilitated rapid cell expansion and established hCD confluence within 3 days (the point of confluency is marked as “day 0” in Fig. 3a and 3d). After rapid increases with proliferation, TEER and TEPD (negative luminal voltage) continued their development, eventually reaching a plateau approximately two weeks after the confluency. However, the subsequent transition to the physiologically advanced media in Phase 2 triggered another dramatic surge in both TEPD and TEER: TEPD doubled, and TEER soared 600-fold beyond Phase 1’s peak. This remarkable shift underscores the profound impact of in vivo biochemical conditions on the functional advancements in tight junctions and ion transport of hCD epithelium.

**Figure 3.**
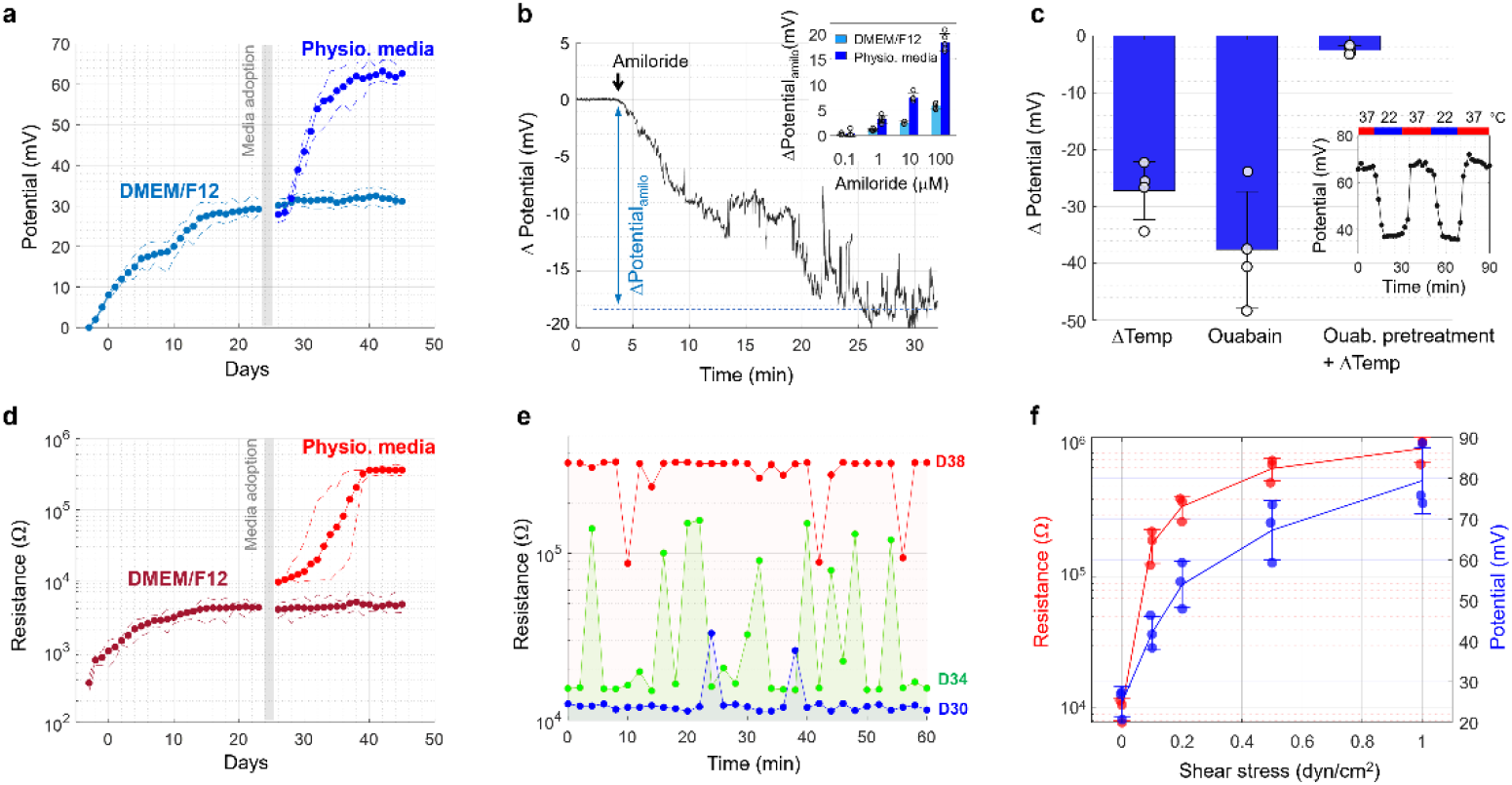
Long-Term Real-Time Measurements of Transepithelial Electrophysiology Reveal Functional Development of hCD in Epi-MAP’s Advanced Physiological Maturation. **a & d**, Exploring dynamics of transepithelial potential difference (TEPD) and resistance (TEER) during two-stage hCD culture in Epi-MAP. In Phase 1, utilizing DMEM/F12-based epithelial media, a full epithelial monolayer confluency is rapidly established (notated as day 0), accompanied by a swift increase in TEPD and TEER. While TEPD and TEER levels gradually stabilize in Phase 1, Phase 2, employing more physiologically relevant media, unveils additional significant surges, emphasizing the crucial role of the biochemical environment in transepithelial physiology. The plot depicts measurement averages using lines with circle symbols, while dotted lines forming envelopes indicate the range of the highest and lowest values recorded during measurements. Flow dynamics were maintained as a baseline for 0.2 dyn·cm^-2^ of apical shear stress and 0.1 mL·min^-1^ of basolateral flow. **b**, Transepithelial potential responses to apical amiloride in hCD Epi-MAP. The temporal trace record at a 4Hz rate presents the dynamic TEPD response to the apical ENaC inhibition with 100 µM amiloride, defining an amiloride-sensitive potential by a stabilized level. The inset highlights the disparity in amiloride-sensitive potential levels between hCD Epi-MAP models cultured in conventional and physiological media, revealing that physiological media promote more mature ENaC function in hCD cells. Error bars represent standard deviation of biological replicates (n=5). **c**, Transepithelial potential response to environmental temperature and basolateral ouabain (a Na^+^-K^+^-ATPase inhibitor) application. The inset displays the time trace of reproducible TEPD response to environmental temperature. This temperature-sensitive TEPD was abolished after 50 µM ouabain treatment (the third case in bar plot), indicating Na^+^-K^+^-ATPase as a temperature-sensitive biological element contributing to TEPD generation in the Epi-MAP culture. Error bars represent the standard deviation of biological replicates (n=4). **e**, Transient dynamics during transepithelial resistance development at day 30, 34, and 38, potentially depicting the transitioning progress of the tight-junction network to a mature, high-resistance state shown in an equivalent circuit model (Supplementary Fig. 2). **f**, Proportional scaling of hCD transepithelial functions under long-term apical shear stress of physiological media. The red and blue lines represent TEER and TEPD values, respectively, of hCD models exposed to various apical shear stress for two weeks, starting with identical baseline characteristics at Phase 1 culture. Error bars depict the standard deviation of biological replicates (n=3).

Our investigation into TEPD generation in matured hCD Epi-MAP identified ENaC and Na^+^-K^+^ ATPase as key players. First, apical amiloride application confirmed ENaC’s electrogenic activity on the lumen-negative transepithelial potential at the apical membrane (Fig. 3b). Epi-MAP’s temporal resolution allowed for a detailed examination of amiloride’s inhibition kinetics, revealing transient fluctuations—possibly arising from ENaC’s rapid turnover—and revealing a stable amiloride-sensitive TEPD approximately 30 minutes after application. The dose-dependent inhibition profile further validated ENaC’s electrogenic role and highlighted the enhanced electrogenicity of ENaC in the condition of physiological media compared to conventional media (see inset of Fig. 3b).

Along with apical electrogenic ENaC, Na^+^-K^+^ ATPase played a vital role in generating the lumen-negative transepithelial potential, residing on the basolateral membrane (Fig. 3c). We unraveled its role by modulating activity through temperature and basolateral ouabain (a Na^+^-K^+^ ATPase inhibitor). Consistent with its established temperature sensitivity of Na^+^-K^+^ ATPase in kidney tubules^28-32^, lowering environmental temperature from 37°C to 22°C (while maintaining CO_2_) triggered a 40% decrease in transepithelial potential (inset, Fig. 3c). Similarly, basolateral ouabain application caused a comparable drop, and no further decrease occurred when temperature was lowered after ouabain. While this clarifies the specific roles of this ion pump in the Epi-MAP’s condition, its temperature-dependent response, along with our characterization on temperature-responsive impedance (Supplementary Fig. 4), affirms the critical need for “in-situ” characterization to truly understand epithelial electrophysiology in action.

The dramatic TEER boost, reaching an order of magnitude of 10^5^ Ω·cm^2^, was accompanied by unique temporal patterns not previously reported (Fig. 3e). The time-resolved trace displayed intermittent occurrences of discrete jumps between two resistance levels during the rapid rise phase, gradually converging to a stable plateau over time. This behavior finds explanation in an equivalent circuit model (Supplementary Fig. 2), where high-level tight junctions are represented in parallel. In the condition where the majority of tight junctions form a high-integrity network, the equivalent total resistance (corresponding to TEER) exhibits discrete jumps, as individual tight-junction elements transition to a mature level. Along with the measured values, these findings convincingly present the formation of high-integrity tight junctions within the physiological dynamics of Epi-MAP.

Further delving into the intricate interplay between physiological fluid dynamics and hCD development, we investigated the impact of prolonged apical shear stress of the biomimetic fluid on hCD transepithelial physiology. While previous studies acknowledged the roles of apical shear stress in epithelial function, they were limited by short-term setups and non-physiological media flow^33-36^. In contrast, our study explored the long-term effects of various levels of apical shear stress (0.01 to 1 dyn·cm−^2^) through a consistent two-week flow of physiological media. All other conditions from the DMEM/F12 stage served as a baseline, allowing us to observe changes directly proportional to the applied apical shear stress (Fig. 3f). Exposure to the highest shear stress (1 dyn·cm−^2^) triggered a remarkable 3-fold increase in transepithelial potential and a 100-fold increase in resistance compared to the lowest shear stress. Intriguingly, at the lowest shear stress, TEER and TEPD resembled those of the static transwell setup (Model 3), suggesting minimal impact of basolateral fluidics. This evidence suggests that chronic, physiologically relevant apical shear stress likely plays a pivotal role in refining key transepithelial attributes, potentially acting as a crucial force for postnatal kidney development, demanding further investigation.

### Human Collecting Duct Epithelium in Epi-MAP Exhibits Comprehensive Functional Responses to Vasopressin and Aldosterone

Dynamic response to hormonal stimulation is crucial for faithfully replicating the kidney’s diverse physiology, modeling disease states, and evaluating potential therapeutic agents. In the collecting duct, vasopressin governs water reabsorption, while aldosterone regulates sodium reabsorption, both critical for maintaining fluid and electrolyte balance in the body. Dysregulation of these signals is implicated to various disorders, making a deeper understanding of their effects essential for designing clinical strategies and identifying novel therapeutic agents^37,38^. The hCD Epi-MAP model possesses the capability to respond to these CD-specific signaling. Alongside biological advances, its unparalleled ability proves invaluable in this pursuit by capturing real-time, time-resolved electrophysiological insights into transepithelial ion/water flux and epithelial integrity.

To decipher the intricate dance between hCD and vasopressin signaling, we employed desmopressin, a clinically relevant analog with V2 receptor (V2R) specificity. Upon V2R signaling, collecting duct PCs orchestrate water permeability through dynamic AQP2 regulation^39^ (Fig. 4a). The established osmolarity gradient along the apicobasal axis then directs water flux, promoting retention and electrolyte reduction in the interstitial compartment. In line with the in vivo scenario, basolateral desmopressin perfusion (10 nM) triggered a rapid and substantial decline in basolateral conductivity within 30 minutes, followed by a gradual reach to equilibrium (Fig. 4c). This pattern likely reflects the initial, non-genetic phase of V2R signaling, involving the translocation of the cytoplasmic AQP2 pool to the apical membrane, followed by transcriptional upregulation of water transporter components^40^. The multimodal measurements revealed additional subtle but noticeable shifts in TEPD (Fig. 4f) and TEER (Fig. 4b), lagging by about an hour behind the initial conductivity response. These changes, captured in real-time with remarkable precision, are consistent with established downstream signaling events, such as cytoskeleton remodeling of tight junctions during AQP2 trafficking and increased ENaC activity^41,42^.

**Figure 4.**
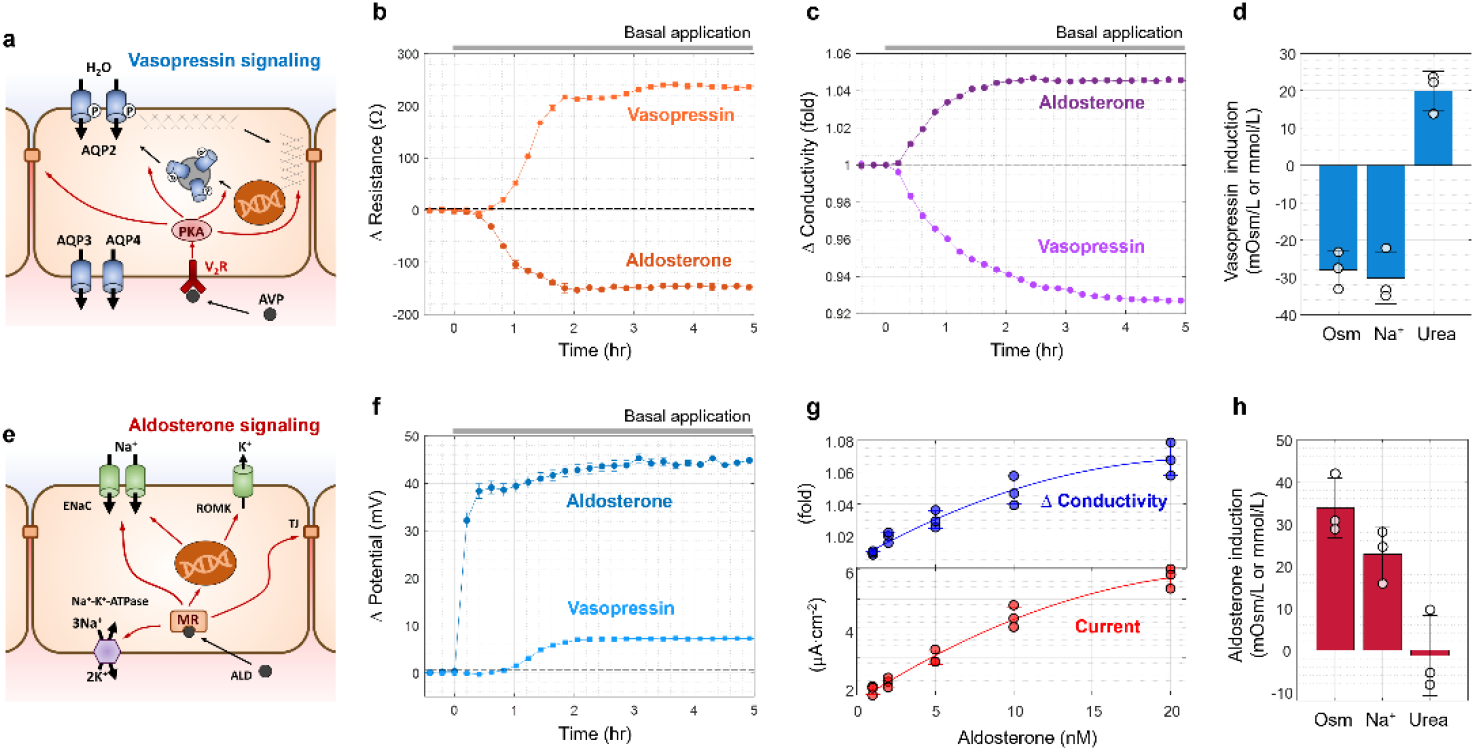
Multimodal Electrophysiology of hCD Epi-MAP: Unveiling Multifaceted hCD Dynamics upon Vasopressin and Aldosterone Signaling. **a**, Illustration of vasopressin signaling in collecting duct principal cells, depicting intracellular changes in aquaporin and cytoskeleton organization. **b, c, & f**, Temporal dynamics of transepithelial resistance, conductivity, and transepithelial potential in response to basal application of desmopressin (a clinically relevant analog with V2 receptor specificity) or aldosterone. In Epi-MAP, each mature hCD epithelium (exhibiting typical transepithelial properties) underwent iterative cycles of the three consecutive measurements, separated by ∼12-minute intervals. Error bars (standard deviation) indicate the spread of measurement values, reflecting short-term variations in each hCD’s physiological expression. **d & h**, Physiochemical analysis of 12-hour interstitial perfusates before and after desmopressin or aldosterone exposure, confirming physiological responses depicted in the electrophysiological readouts. Osmolarity (Osm) is presented in mOsm/L, and sodium (Na^+^) and urea are displayed in mmol/L. **e**, Illustration of aldosterone signaling in collecting duct principal cells, highlighting cellular components affected in the signaling, including sodium, potassium, and tight-junction fluxes. **g**, Dose-dependent response of the collecting duct Epi-MAP model to aldosterone concentrations. Values represent the 3-hour post-aldosterone state. Current is calculated using Ohm’s law of transepithelial potential and resistance. Error bars in d, h, and g depict the standard deviation of biological replicates (n=3).

Post-hoc physiochemical analysis of the 12-hour interstitial perfusate decisively confirmed the desmopressin-induced water transport. Compared to pre-treatment collections, we observed a 28.3 ± 4.9 mOsm·L−^1^ decrease in osmolarity, a 30.2 ± 7.4 mM decrease in sodium, and a 20.1 ± 4.8 mM increase in urea (Fig. 4d). These changes point to V2R signaling’s key role in water transport: the reduced sodium concentration reflects the larger water influx diluting the overall electrolyte concentration, while the urea increase is consistent with V2R-mediated urea permeability as another water transport mechanism. Leveraging our comprehensive quantitative measurements and controlled parameters, we further delved into these dynamics with a computational Multiphysics simulation (details in Method). Enabled by Epi-MAP’s precise design, this approach revealed a net water flux of 50nl·min^-1^·cm^-2^, or a hydraulic conductivity (*L*_*p*_ = *J* · *A*^−1^ · ΔΦ^−1^, where *J* is water flux, *A* is an epithelial area, and ΔΦ is osmotic pressure difference) of 5E-6 cm·atm^-1^ ·sec^-1^, aligning with the expected range observed in experiments with isolated perfused tubules^43-45^. This alignment strongly suggests in vivo-like functional maturation of hCD in the Epi-MAP culture, solidifying its potential to simulate hCD-specific physiological responses with diverse biochemical substances.

Similarly, aldosterone stimulation on the hCD model exhibited multi-parametric dynamics in the electrophysiological traces, reflecting its in vivo functions on the human collecting duct. As aldosterone acts primarily on PCs via the mineralocorticoid receptor to enhance ENaC and Na^+^-K^+^ ATPase activity (Fig. 4e), the major response was observed in TEPD, showing a rapid increase within the first 20 minutes, followed by a gradual rise over the next several hours. These dynamics reflect both rapid non-genomic action and long-term transcriptional changes observed in vivo^46,47^. The increased sodium transport was evident in the gradual rise of interstitial conductivity (Fig. 4c) and an increase in osmolarity and sodium concentration in the offline physiochemical assay (Fig. 4h). The comprehensive transepithelial physiology was shown with a high-fidelity dose-responsiveness (Fig. 4g), while excluding untargeted responses such as urea transport (Fig. 4h). Additionally, a minor decrease in transepithelial resistance (Fig. 4b) hinted at aldosterone’s influence on paracellular conductivity, aligning with prior studies on claudin phosphorylation^48^. Notably, our hCD Epi-MAP model responded to basolateral, but not apical, aldosterone application, despite its lipophilic nature^49^, revealing the maturation of hCD apical membrane with molecular uptake selectivity.

## DISCUSSION

This study introduces an innovative Epi-MAP system, precisely designed to replicate the complex functional architecture of the human kidney collecting duct (hCD). Utilizing hCD cells derived from human pluripotent stem cells, we have established a reproducible and physiologically advanced hCD epithelium within a precisely defined biomimetic environment. This integrated platform boasts continuous, in-situ, and multi-scale electrophysiological measurements, enabling us to decipher the intricate influence of microenvironmental cues on hCD functional maturation. The comprehensive physiological responses of the hCD epithelium, characterized through seamless monitoring of Epi-MAP, solidifies this modeling approach as a powerful tool for translating ex vivo differentiation into an experimentally advanced in vitro model. This model unlocks new avenues for understanding the intricate physiological complexities of kidney tubule function and pathology.

Firmly anchored in established knowledge of collecting duct physiology, Epi-MAP unravels the intriguing complexities of hCD development, illuminating previously hidden facets of CD physiology. Its meticulous examination exposes captivating characteristics, including the spontaneous emergence of interlocking principal and intercalated cell arrangements within a dynamic physiological milieu; surpassing current estimations of tight junction integrity; and prolonged apical fluidic flow functioning as a crucial regulator of dynamic transepithelial physiology. Furthermore, the quantitative, comprehensive analytics of Epi-MAP provides a powerful spectrum of multifaceted responses, paving the way for their utilization as an innovative drug discovery platform. Lastly, our study underlines the critical value of “in situ” monitoring, not only for ensuring data fidelity but also for safeguarding physiological relevance, offering a valuable insight with significant implications for future research.

Envisioned future directions for Epi-MAP include its readily attainable scalable production to enable high-throughput investigations into diverse physiological and pathological states, propelling pharmaceutical development and therapeutic discovery. Additionally, expanding our access to robust and reliable hiPSC-derived physiological cell sources will bolster our pathophysiological investigations by unlocking disease-specific genetic signatures. This concurrent development of a sophisticated platform and diverse cell sources exemplifies a powerful approach to enhance drug target identification and validation in a more reproducible and measurable manner.

While Epi-MAP has successfully elucidated key aspects of human kidney function, its potential is further amplified by addressing current limitations. One constraint relates to its current measurement of bulk transepithelial ion transport, restricting our capacity for detailed investigation of specific ion movements across the collecting duct and other epithelia. To address this, we are actively developing the integration of microelectrode-based ion flux measurement, which will provide the precision necessary to decipher the intricate interplay of individual ions in human kidney and other organ physiology and pathophysiology. Another limitation is the lack of cellular proliferation simulation within the interstitial fluid, a driving force behind many acquired and genetic kidney diseases affecting the nephrovascular unit. Recognizing this, we are exploring the incorporation of natural tissue-derived membranes alternative to the current polymer nanoporous membrane, which can be remodeled by cellular activities. We anticipate that these modifications will enable the modeling of intricate tubular morphogenesis in disease states, providing us with the capability to more comprehensively dissect human kidney pathology.

Epi-MAP represents a revolutionary shift in human physiological modeling, catalyzing the emergence of a future where individual patient physiologies guide treatment decisions. By embracing and strategically overcoming current limitations, we aim to harness the transformative power of this platform, paving the way for therapeutic interventions directly dictated by the unique language of each patient. Epi-MAP is not just a technological marvel, but a groundbreaking translation tool for bridging the research-to-clinic gap, illuminating the path towards a future where the patient’s own biology informs therapeutic decisions. While the road to personalized kidney care remains complex, Epi-MAP offers a powerful tool to accelerate our understanding of human kidney physiological diversity and disease mechanisms, paving the way for more informed and effective therapeutic approaches in the future.

## METHODS

### Fabrication of integrated Epi-MAP microfluidic device

The microfluidic device was an assembly of two polymer fluidic channels, one nanoporous membrane (PETE membrane, 0.4-micron pores, 2E6 pores/cm^2^, Steritech), an electrode-patterned glass substrate, and two media reservoirs. The microfluidic channels were prepared as PDMS (DOW SYLGARD™ 184) replication from a SU8 mold fabricated by UV photolithography using SU8 3035 (Kayaku Advanced Materials) following the product manual. In separate fabrication, the surface electrode was patterned on a glass slide by a lift-off process using LOR3A, SU8 3005, and titanium/platinum (5/50nm) evaporation (ROCKY Mountain Vacuum Tech). The electrode material was chosen based on chemical stability, interfacial impedance, and electrical noise (Supplementary Fig. 3). For the assembly, first, the nanoporous membrane was modified with a 15-minute incubation of 1% (3-Aminopropyl)triethoxysilane (APTES, Sigma-Aldrich) and sandwiched between oxygen plasma-treated PDMS chips. The hole-drilled electrode-patterned glass was bonded with the PDMS assembly through oxygen plasma and attached with media reservoirs fabricated by CO2 laser machining using uncured PDMS. Sterilization before cell loading was achieved through ultraviolet ozone (UVO) treatment and overnight incubation of PBS with 1% penicillin streptomycin (ThermoFisher Scientific).

### Automated fluidic dynamics and electrophysiological monitoring

To minimize experimental hassles of microfluidic operation during long-term maturation of the epithelial model, we designed the continuous perfusion of Epi-MAP using a home-built tilting plate system, changing the angular position of a large plane plate (40cm x 40cm) according to a scheduled program in a microprocessor (Arduino Uno) with an angular sensor (ADXL345, Analog Devices). The combination of tilting rate, reservoir size, and inter-inlet distance was determined for specific flow rates of luminal and basal channels (2 µl·min^-1^ and 0.1 µl·min^-1^ unless indicated), minimizing device intervention (media replenishment every two days). To enhance flow rate precision during the conductivity measurement, we used a two-channel syringe pump with 1mL and 20mL syringes. The ratio of inside diameters of the two syringes generated an 18.2-fold difference in flow rates, that was 2 µl·min^-1^ and 0.11 µl·min^-1^ in the luminal and basal channels, respectively.

For the electrical recording, the contact pads along a glass slide edge were wire-connected to a multichannel relay. The computer-programmed switching of the relay was designed to connect two-point terminals of an electrochemical station (CHI750A, CH Instruments, Inc.), autonomously synchronized for serial measurement of AC impedance and open circuit potential. To automatically link the relay switching and operation of the electrochemical station, we used a BASIC-like scripting language, AutoIt (Ver. 3) and CHI750A script (CH Instruments, Inc.).

### Human collecting duct epithelial cells

The preparation of human collecting duct epithelial cells is described in our earlier study^24^ using H9 hESC-derived kidney organoids. Then, this study was designed using a large batch of cryopreserved epithelial cells at passage 15. After rapidly thawing the frozen cells in a 37 ?C warm bath, the cells in the cryopreservation reagent (10% DMSO addition to the culture media) were transferred to prewarm DMEM/F12 in a 15 ml tube and centrifuged at 980rpm to leave a cell pellet. The cells were gently resuspended and seeded into a 6-well plate with DMEMological models, potentially limiting its applicabilit% v/v, ThermoFisher Scientific), epithermal growth factor (10 ng/mL, ThermoFisher Scientific), T3 (1 nM, Tocris) and fetal bovine serum (2% v/v, ThermoFisher Scientific). The cells at 80-90% confluency were dissociated into single cells using Trypsin-EDTA and seeded into the study models including 12-well plates, transwell inserts (Corning), and the apical channel of Epi-MAP. The single-cell concentration was adjusted for approximately 30% surface coverage at the initial attachment. The static culture models (Model 1, 2, and 3) had media changes every 2 days. In the dynamic culture in Epi-MAP (Model 4), the designed media perfusion rates began one day after the loading. The media transition to physiological media was implemented by gradually increasing the percentage of physiological media by 25% per day for four days.

### Physiological media for human collecting duct epithelial cells

A set of physiological media were prepared as following: For the luminal fluid, the media was composed of 4 mM creatinine, 5 mM Na_3_C_6_H_5_O_7_, 30 mM KCl, 15 mM NH_4_Cl, 3 mM CaCl_2_, 2 mM MgSO_4_, 2 mM NaHCO_3_, 0.1 mM NaC_2_O_4_, 9 mM Na_2_SO_4_, 3.6 mM NaH_2_PO, 0.4 mM Na_2_HPO_4_ with additional NaCl for the final concentration of Na^+^ to be 100mM and 200mM urea (all components were purchased from Millipore Sigma). For the interstitial fluid, DMEM/F12 media was based with 2% fetal bovine serum, 1% Insulin-Transferrin-Selenium, 1nM T3, 100mM urea, and additional NaCl for Na^+^ to be 200mM. The final osmolarities of each medium were 460 and 480 mOsm·L^-1^, respectively. Detailed information is provided in Supplementary Table 1.

### Analysis of epithelial morphology

Epithelial monolayer morphologies were imaged using a 10X objective and phase-contrast annulus on an inverted microscope (CKX41, Olympus). Subsequently, these images were subjected to analysis using the ImageJ Fiji software with a designated plug-in app.

The app’s settings were empirically optimized to automatically define regions of interest (ROI) based on cell boundaries. It also recorded the individual area and aspect ratio (AR) of these ROIs while generating pseudocolor images with analytical results. To characterize the underlying patterns and characteristics of epithelial morphology, a statistical distribution was presented in histograms. During the analyses, regions containing ambiguous cellular boundaries and instances of multi-layered growth were meticulously excluded through manual intervention to ensure the accuracy of the findings.

### Immunostaining and transcriptomic analysis

Immunostaining of cells in Epi-MAP was conducted with serial perfusions of following reagents sequentially through the luminal channel: (1) 4% paraformaldehyde for one hour; (2) PBS wash for one hour; (3) 0.1% triton-X permeabilization for one hour; (4) 2% BSA PBST blocking for eight hours, (5) primary antibody staining in 1% BSA PBST for eight hours; (6) 1% BSA PBST wash for eight hours; (7) secondary antibody and DAPI staining in 1% BSA PBST for eight hours; and (8) PBST wash for eight hours. PBST was composed of a 0.2-micron syringe filtered solution of 0.1% tween-20 in PBS. The primary antibodies used in this study included anti-SCNN1A (Millipore Sigma, HPA012743), anti-Aquaporin 2 (Abcam, ab199975), anti-Acetyl-α-Tubulin (Cell Signaling, 5335), and anti-V-ATPase B1/2 (Santa Cruz Biotechnology, sc-55544) and were prepared as dilutions of 1:200, 1:100, 1:500, and 1:50, respectively. Secondary antibodies were used at a 1:500 dilution to match the target species (Millipore Sigma). All the steps were conducted under low-light conditions at room temperature. Immunostaining was imaged using an inverted laser scanning confocal microscope (Zeiss LMS980) and reconstructed as a z-axis projection using ImageJ (National Institution of Health)

RNeasy Plus Mini Kit (Qiagen) was used for on-chip lysis and RNA purification. The process involved perfusing a lysis buffer (Buffer RLT Plus) through the luminal channel to lyse the cells on the membrane, followed by retrieval of lysates from the chip by pipetting at the outlet. Following the manufacturer’s protocol, RNA was purified from the lysates and immediately converted into complementary DNA (cDNA) using a reverse transcription kit (iScript Reverse Transcription Supermix, Bio-Rad). The resulting cDNA samples were stored at -80°C until PCR analysis. Three biological replicates were prepared from separate experiments for statistical analysis.

Real-time PCR analysis was performed using the SsoAdvanced Universal SYBR Green Supermix (Bio-Rad) and the CFX96 real-time PCR system (Bio-Rad). The PCR cycle consisted of 40 cycles with denaturation at 95°C and annealing at 60°C. PCR byproduct amplicons were verified through melting analysis (65-95°C in 0.5°C increments). Primers used in this study were purchased from Predesigned qPCR Assays (Integrated DNA Technologies) or qPCR Primer Pairs (OriGene Technologies, Inc.).

### Computational simulation

We conducted a comprehensive analysis to estimate the transepithelial flux of water and sodium, combining computational simulations of a simplified channel model for the Epi-MAP interstitial channel with post-hoc physiochemical analysis of collected perfusates.

Using COMSOL Multiphysics (ver. 5.2), we constructed a straight, rectangular cross-sectional channel with geometric properties mimicking those of the Epi-MAP’s interstitial channel. We performed simulations incorporating fluidic dynamics, molecular transport, and electric conductivity of the conductive medium. The inlet and outlet boundaries were configured to simulate laminar inflow with a constant sodium concentration input, which was experimentally determined. We simplified the epithelial layer by introducing a boundary condition that generated a uniform flux of both water and sodium. We determined changes in electrical conductivity, primarily assuming that conductivity is largely dependent on sodium concentration. The sodium flux across the epithelium was presumed to vary proportionally with changes in transepithelial potential.

Through a systematic iterative process, we adjusted the simulation parameters for sodium and water flux, subsequently comparing the simulated results with experimental measurements of sodium concentration at the outlet. This comparative analysis allowed us to estimate transepithelial fluxes.

### Characterization of perfusates

We analyzed the 12-hour perfusate collections from the outlet of the interstitial channel for sodium concentration, urea concentration, and osmolarity. Sodium concentration was quantified using a fluorescent indicator called ING-2 TMA+ from Ion Biosciences, following the manufacturer’s instructions. Urea concentration was determined using a colorimetric array LS-K331 from LS Bio, also following the manufacturer’s recommended protocol. Osmolarity was measured using a vapor pressure osmometer (Wescor 550). To enhance calibration accuracy, we performed an intermediate calibration using a fresh calibration ampule after every six measurements. This meticulous calibration procedure was employed to ensure the precision of our measurements from ambient temperature drift. To ensure the accuracy of our measurements, all samples were subjected to analysis in technical triplicates.

## Acknowledgments

This work is supported by grant no. NIH NCAT 5UH3TR002155-05. J.V.B. was also supported by NIH R37 DK39773 and DK 072381. We thank J. Cho for technical assistance.

## Author Contributions

L.P.L. and J.V.B. proposed and supervised the project. S.H., M.S., J.V.B., and L.P.L. designed the study. S.H. and M.S. performed experiments for data. A.P. and K.M. generated ureteric bud organoids from human embryonic stem cells and an epithelial cell. S.H. wrote the manuscript. All authors discussed and edited the manuscript.

## Data Availability

All the data related to this study have been comprehensively presented in the manuscript, Supplementary Information, and Source Data file. Any further data can be available upon reasonable request from the corresponding author.

## Notes

### Competing Interest Statement

J.V.B. and K.W.M. are co-inventors on kidney organoid patents assigned to Mass General Brigham.

