## Supplementary Information for "Human Stem Cell-derived Kidney Collecting Duct Model via Epithelial Microphysiological Analysis Platform: Epi-MAP"

- *Contents*

  - *Supplementary Figure 1: Impact of Fluidic Shear Stress on Ciliary Expression*
  - *Supplementary Figure 2: Estimation of transepithelial resistance induced by localized tight-junction degradation.*
  - *Supplementary Figure 3: Optimization of surface electrode integration and material.*
  - *Supplementary Figure 4: Impact of Environmental Temperature on Resistance Measurement in MAP.*
  - *Supplementary Figure 5: Electrical conductivity response to additional sodium chloride concentration.*
  - *Supplementary Table 1: Components of physiological media used in hCD Epi-MAP culture.*

**Supplementary Figure 1. Impact of Fluidic Shear Stress on Ciliary Expression.** Staining with anti-acetyl-α-tubulin antibody underscores the influence of fluidic shear stress on the ciliary expression of hCD cultured in physiological media. **a,** Comparison of cilia length between Model 3 and Model 4. **b,** Comparison of the percentage of ciliated cells in the epithelial monolayer of Model 3 and Model 4. **c,** A representative image of acetyl-α-tubulin immunostaining in Model 4.

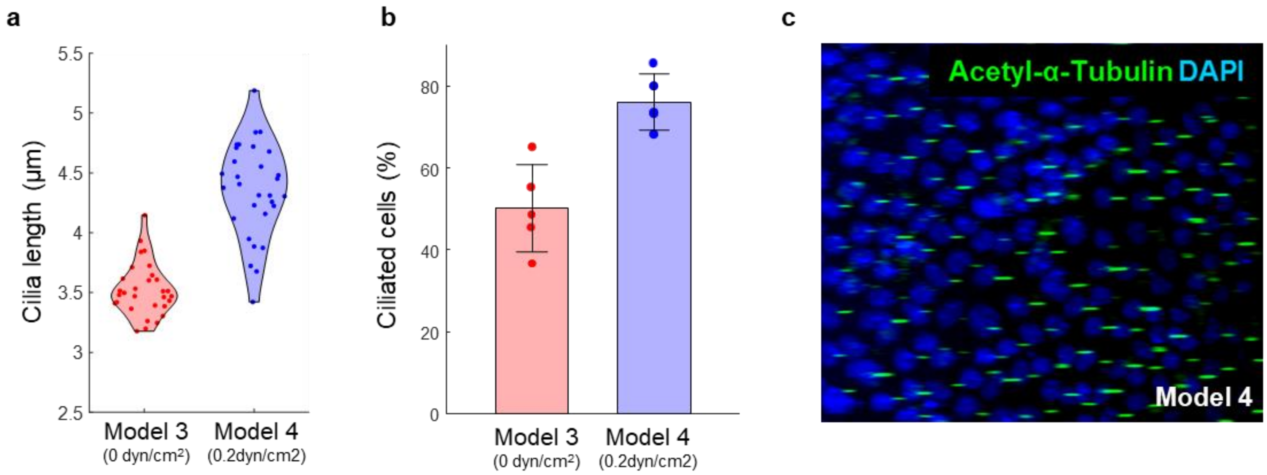

**Supplementary Figure 2. Estimation of transepithelial resistance induced by localized tight-junction degradation. a,** Schematic representation of the configuration for measuring transepithelial electrical resistance across an epithelial monolayer. **b,** Equivalent electrical model illustrating transepithelial electrical resistance. **c,** Normalized transepithelial resistance calculated across the proportion of weakened tight junctions, varying the ratio of "good" and "bad" tight-junction equivalent resistance. **d,** Detailed view of the calculated normalized transepithelial resistance. This finding suggests a possible explanation for the discrete jumps observed in the time-resolved TEER measurements in Figure 3e. These jumps may indicate the progressive incorporation of high-resistance tight junctions into a well-established tight-junction network, with a small portion potentially remaining in a less developed lower-resistance state.

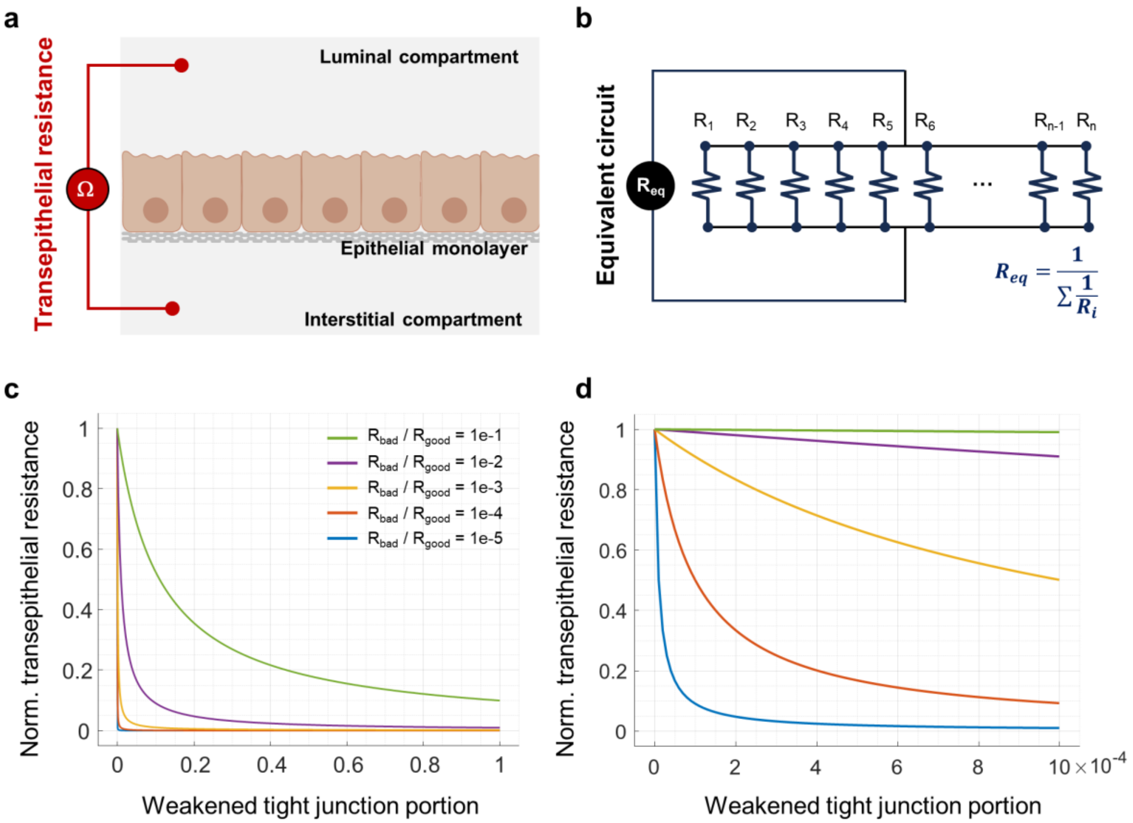

**Supplementary Figure 3. Optimization of surface electrode integration scheme and fabrication material. a,** Two possible configurations of surface electrode integration within the Epi-MAP fluidic chip are depicted. The upper illustration shows the proximal electrode integration used in this study, while the lower one demonstrates a configuration with surface electrodes positioned at the fluidic ports. The first configuration exhibited a six-fold lower resistance in the blank Epi-MAP without cells, attributed to the proximity between the electrode sets. **b,** In the Epi-MAP model's proximal electrode integration, the shown blank resistance measurements reveal the outstanding performance of the pre-fixed electrode-electrolyte interface. This design delivers exceptional signal-to-noise ratios exceeding 2000, ensuring temporal stability and reliable electrical readouts. **c & d,** Characterization of surface electrode materials with respect to their electrical resistance, noise level, and chemical stability. While standard silver/silver chloride electrodes (1.5 mm diameter) achieved the lowest resistance, their high noise, likely due to interfacial reactions, led us to choose platinum. This metal exhibits moderate resistance, excellent noise characteristics, and notable chemical stability. These analyses were based on measurements across two electrodes placed on a 3 cm long, 0.5 mm wide channel for comparison with silver/silver chloride wire.

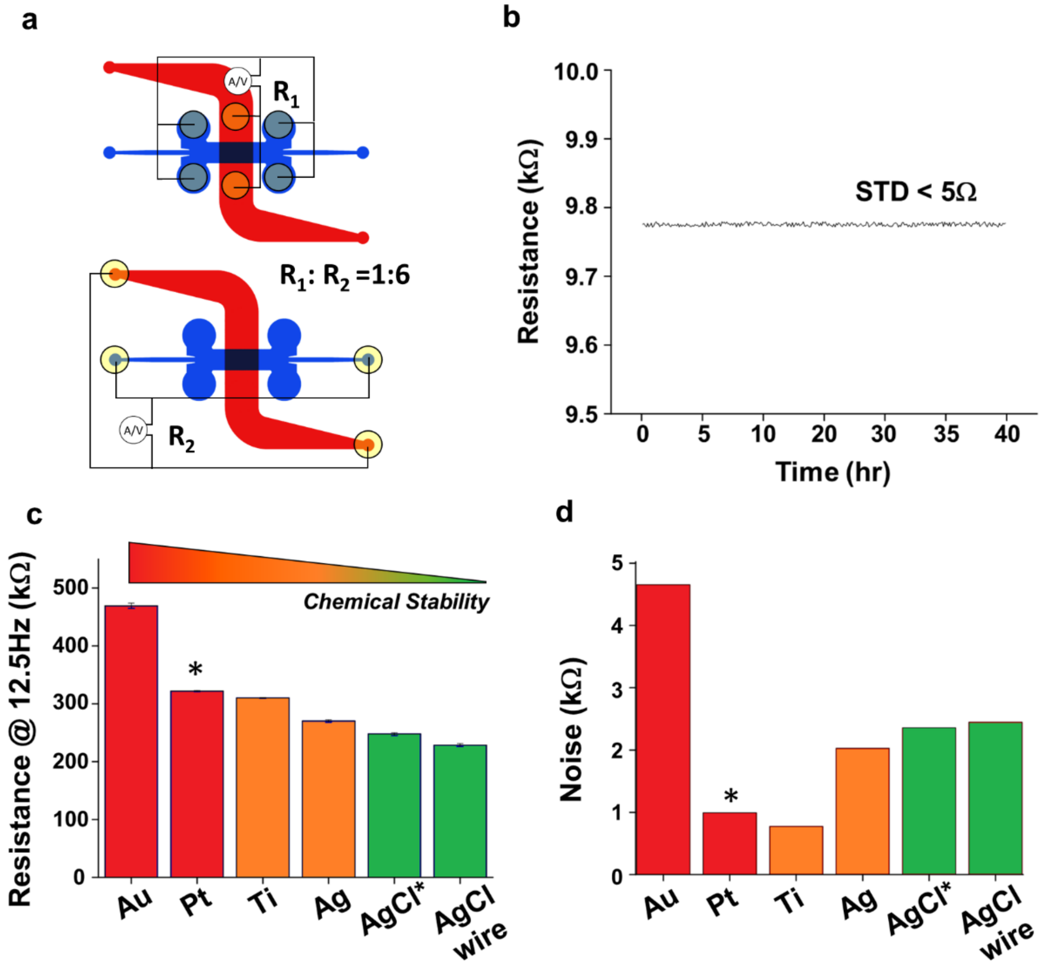


(*) E-beam deposition of AgCl on Ag/glass electrode

**Supplementary Figure 4. Impact of Environmental Temperature on Transepithelial Resistance Measurements in Cell-Free Epi-MAP.** An Epi-MAP device underwent a characterization of background electrical resistance while varying the environmental temperature, both with and without CO2 changes. The observed deflection in electrical resistance due to environmental temperature underscores the importance of in-situ electrophysiological measurement, even without cells. As anticipated from the perspective of heat transfer, the most substantial changes were observed at the initiation of the temperature change followed by a one-hour-long gradual transition.

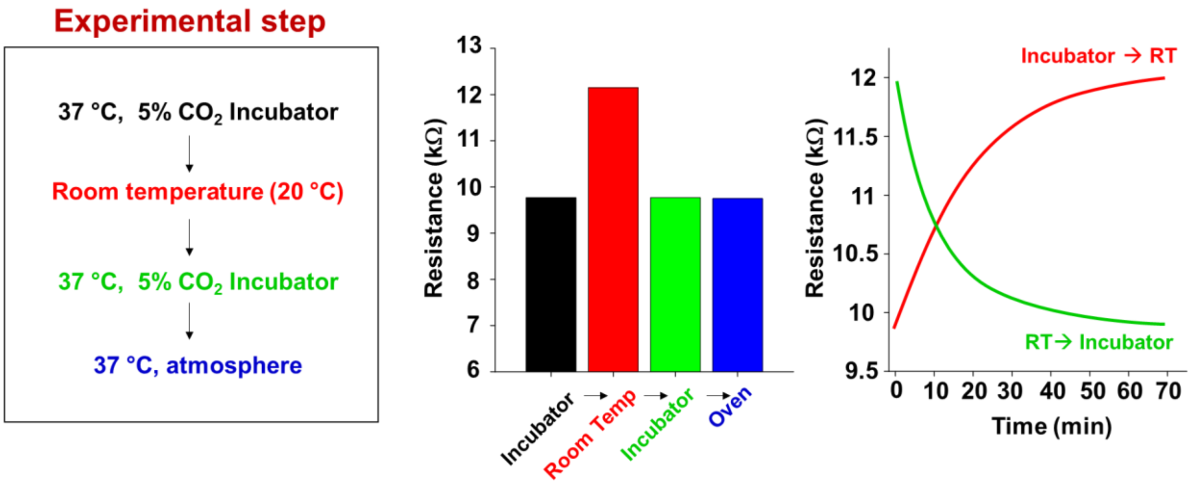

**Supplementary Figure 5. Electrical Conductivity Changes in Physiological Interstitial Media as a Function of Sodium Concentration.** As the most abundant ion among the media components, the addition of sodium chlorides governed the electrical conductivity with a highly linear relationship. This measurement was conducted with fresh interstitial media by gradually adding sodium chloride in the range of 0 to 90. These results were then utilized in the computational simulation to estimate transepithelial sodium flux.

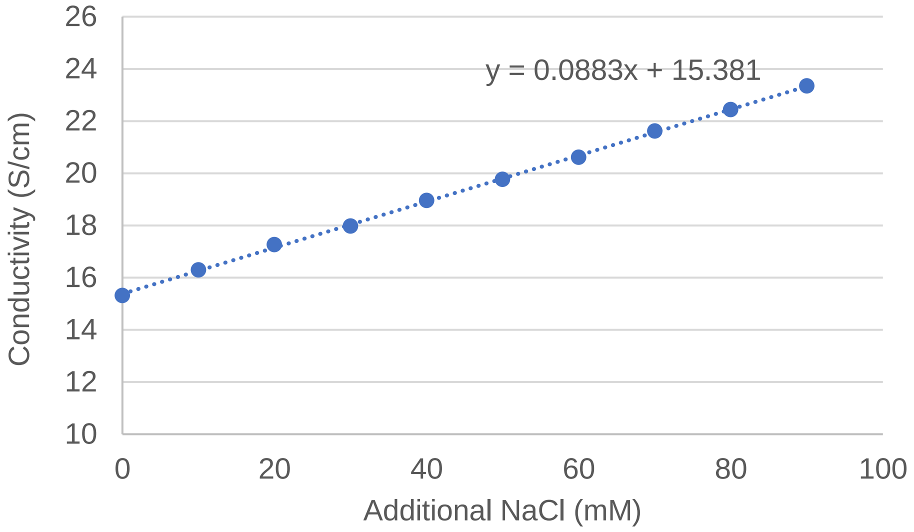

**Supplementary Table 1. Components of physiological media used in hCD Epi-MAP culture.** The bioreagent-grade components listed in the table were prepared as stocks, each nearly at its maximum aqueous solubility. The diluted mixture was then prepared with 0.2-μm filtration, in a bulk volume for a week's usage. The basal media were freshly prepared for a week's usage with the addition of FBS, ITS, T3, and prefiltered (to prevent contamination) sodium chloride and urea.

| Apical media | | |
|---|---|---|
| **Component** | **Concentration** | **Formula** |
| Urea | 200 mM | CO(NH2)2 |
| Creatine | 4 mM | CNCH2CO2H |
| Sodium Citrate | 5 mM | Na3C6H5O7·2H2O |
| Sodium chloride | 60.5 mM | NaCl |
| Potassium chloride | 30 mM | KCl |
| Ammonium chloride | 15 mM | NH4Cl |
| Calcium chloride | 3 mM | CaCl2·2H2O |
| magnesium sulfate | 2 mM | MgSO4·7H2O |
| Sodium bicarbonate | 2 mM | NaHCO3 |
| Sodium oxalate | 0.1 mM | NaC2O4 |
| Sodium Sulfate | 9 mM | Na2SO4 |
| Monosodium phosphate | 3.6 mM | NaH2PO4·H2O |
| Disodium phosphate | 0.4 mM | Na2HPO4 |

| Basal media | | |
|---|---|---|
| **Component** | **Concentration** | **Formula** |
| DMEM/F12 | basal media | |
| Fetal bovine serum | 2% | |
| Insulin-Transferrin-Selenium | 1% | |
| Triiodothyronine | 1nM | C15H12I3NO4 |
| Sodium chloride | 60mM | NaCl |
| Urea | 100mM | CO(NH2)2 |
